# Contrasting macroevolutionary patterns in the human N-glycosylation pathway

**DOI:** 10.1101/2024.10.30.621208

**Authors:** Domagoj Kifer, Nina Čorak, Mirjana Domazet-Lošo, Niko Kasalo, Gordan Lauc, Göran Klobučar, Tomislav Domazet-Lošo

**Affiliations:** Faculty of Pharmacy and Biochemistry, University of Zagreb, A. Kovačića 1, HR-10000 Zagreb, Croatia; Laboratory of Evolutionary Genetics, Division of Molecular Biology, Ruđer Bošković Institute, Bijenička cesta 54, HR-10000 Zagreb, Croatia; Department of Applied Computing, Faculty of Electrical Engineering and Computing, University of Zagreb, Unska 3, HR-10000 Zagreb, Croatia; Genos Glycoscience Research Laboratory, Borongajska cesta 83h, HR-10000 Zagreb, Croatia; Department of Biology, Faculty of Science, Division of Zoology, University of Zagreb, Rooseveltov trg 6, HR-10000 Zagreb, Croatia; School of Medicine, Catholic University of Croatia, Ilica 242, HR-10000 Zagreb, Croatia

## Abstract

Building on coding mutations and splicing variants, post-translational modifications add a final layer to protein diversity that operates at developmental and physiological timescales. Although protein glycosylation is one of the most common post-translational modifications, its evolutionary origin remains largely unexplored. Here, we performed a phylostratigraphic tracking of glycosylation machinery genes and their targets — glycosylated proteins — in a broad phylogenetic context. Our results show that the vast majority of human glycosylation machinery genes trace back to two evolutionary periods: the origin of all cellular organisms and the origin of all eukaryotes. This indicates that protein glycosylation is an ancient process likely common to all life, further elaborated in early eukaryotes. In contrast, human glycoproteins exhibited prominent enrichment signals in more recent evolutionary periods, suggesting an important role in the transition from metazoans to vertebrates. Focusing specifically on the N-glycosylation pathway, we noted that the majority of N-glycosylation genes acting on the cytoplasmic side of the endoplasmic reticulum (ER) trace back to the origin of cellular organisms. This sharply contrasts with the rest of the N-glycosylation pathway, which is oriented toward the ER lumen, where genes of eukaryotic origin predominate. In the Golgi, we also identified an analogous binary evolutionary origin of glycosylation machinery genes. We discuss these findings in the context of the evolutionary emergence of the eukaryotic endomembrane system and propose that the ER evolved through the invagination of a prokaryotic cell membrane containing an N-glycosylation pathway.

## Introduction

Glycosylation is the enzymatic process that transfers an oligosaccharide (glycan) to an aglycon; i.e., protein, lipid or RNA (Flynn et al. 2021). In *Homo sapiens*, glycans are synthesized and attached to aglycons through one of the 16 glycosylation pathways consisting of about 700 genes encoding enzymes transporters, and other proteins found in cellular glycosylation machinery (Cantarel et al. 2009, Moremen et al. 2012, Varki et al. 2015, Schjoldager et al. 2020). Protein glycosylation is a complex post-translational modification occurring almost completely in the endoplasmic reticulum (ER) and the Golgi (Colley et al. 2017), where glycans are attached to proteins in a series of reactions catalyzed by glycotransferases and glycoside hydrolases (Stanley et al. 2017). It has been known for a long time that glycans play major metabolic, structural and physiological roles in biological systems and that more than a half of all human proteins are glycosylated (Apweiler et al. 1999; Lauc et al. 2014, Varki 2017).

Protein glycosylation can be found in all three domains of life (Chung et al. 2017, Joshi et al. 2018). Compared to bacteria and archaea, eukaryotes use a narrower spectrum of monosaccharide building blocks in glycan synthesis (Springer and Gagneux 2016, Gagneux et al. 2017). However, eukaryotic glycans, despite the reduced panel of monosaccharide units, show highly complex linear and branching structural diversity (Bishop and Gagneux 2007, Lauc et al. 2014, Springer and Gagneux 2016, West et al. 2021). Protein glycosylation can be viewed as a polygenic trait, where glycosylation phenotypes result from the actions of many genes along the glycosylation pathways, partly modulated by intracellular and extracellular environments (Bishop and Gagneux 2007, Suzuki 2019). The studies of protein-bound glycan diversity in different evolutionary lineages showed remarkably discontinuous patterns that are shaped by various evolutionary forces, including coevolutionary interactions within a specific holobiont (Moran et al. 2011, Bishop and Gagneux 2007, Corfield and Berry 2015, Schröder and Bosch 2016, Suzuki 2019, West et al. 2021). However, despite the biological importance and ubiquitous presence of glycosylation on the tree of life (Corfield and Berry 2015, Gagneux et al. 2017; Chung et al. 2017), our understanding of the evolutionary dynamics of glycosylation machinery (GM) genes is still cursory (Tomono et al. 2015, Lombard 2016b, Wang et al. 2017b, Petit et al. 2018, Nikolayev et al. 2020).

For instance, N-glycosylation biosynthesis is the most frequent and the most studied type of protein glycosylation in eukaryotes (Apweiler et al. 1999, Lombard 2016). It starts at the cytoplasmic side of the ER membrane where monosaccharides are sequentially added by glycosylation machinery proteins to the activated lipid carrier until a glycan with Man5GlcNAc2 structure is formed (Stanley et al. 2017). This glycan structure is then translocated by a flippase to the luminal side of the ER membrane, where it is further elongated by the luminal GM to the Glc3Man9GlcNAc2 structure, which is common to most eukaryotes (Corfield and Berry 2015). Further glycan editing continues in the Golgi where, under the control of Golgi-specific GM proteins (glycosyltransferases and glycosidases), their final structure is formed (Stanley et al. 2017). In eukaryotes this final glycan structure varies within populations and between species (Corfield and Berry 2015, Gagneux et al. 2017; Chung et al. 2017; Wang et al. 2017, West et al. 2021).

Eukaryotic, bacterial and archaeal N-glycosylation pathways share a common topology where the first part of stepwise oligosaccharide synthesis unfolds on the cytosolic side of a membrane. At some point this process includes flipping of the nascent oligosaccharide across the membrane (Nothaft and Szymanski 2010, Lombard 2016b, Chung et al. 2017, Eichler and Imperiali 2018). In addition, N-glycosylation pathways show chemical resemblance that include the use of polyisoprenol lipid carriers and phosphate-linked substrates (Jones et al. 2009, Lombard 2016a, Eichler and Imperiali 2018). These topological and chemical similarities suggest that the last universal common ancestor (LUCA) possessed an N-glycosylation-like pathway (Lombard 2016a). The remarkable difference, however, is that in bacteria and archaea, the N-glycosylation pathway is embedded in the plasma membrane, whereas in eukaryotes, it is part of the ER membrane (Nothaft and Szymanski 2010, Corfield and Berry 2015, Lombard 2016b, Li et al. 2017, Chung et al. 2017, Eichler and Imperiali 2018).

On the other hand, the evolutionary origin of individual glycosylation machinery genes acting in the eukaryotic N-glycosylation pathway is not yet fully resolved (Lombard 2016b). Because of the symbiotic origin of eukaryotes with the contribution of both archaeal and bacterial ancestors, it is not fully clear which prokaryotic group is ancestral to the eukaryotic N-glycosylation pathway (Baum and Baum 2014, Gould et al. 2016, López-García and Moreira 2020). Interestingly, recent phylogenomic analysis suggests a mixed origin of eukaryotic N-glycosylation with the contribution of both archaea and bacteria (Lombard 2016b). Nevertheless, the evolutionary origin of many eukaryotic N-glycosylation genes remains unresolved and some of them seem to be a eukaryotic innovation (Lombard 2016b).

Another facet of the glycosylation process involves proteins that are targeted by the glycosylation machinery. At least in humans, it is clear that many proteins are glycosylated, however evolutionary dynamics of these glycoproteins (GPs) has not been addressed so far. To obtain a coherent view of the evolutionary origin of glycosylation machinery and glycoprotein genes, and consequently to gain a better understanding of the role of glycosylation in the evolution of metazoans, particularly humans, we applied here a phylostratigraphic approach (Domazet-Lošo et al. 2007, Domazet-Lošo and Tautz 2008, Domazet-Lošo and Tautz 2010a, Domazet-Lošo and Tautz 2010b, Tautz and Domazet-Lošo 2011, Šestak et al. 2013, Šestak et al. 2015, Domazet-Lošo et al. 2017, Shi et al. 2020, Futo et al. 2021, Čorak et al. 2023, Domazet-Lošo et al. 2024). Our results showed that glycosylation machinery and glycoprotein genes have divergent macroevolutionary dynamics and that the intracellular localization of N-glycosylation proteins on the endoplasmic reticulum membrane follows an evolutionarily polarized pattern. This finding led us to propose that the endoplasmic reticulum evolved through the invagination of the prokaryotic plasma membrane.

## Results

To reconstruct the evolutionary origin of *H. sapiens* glycosylation machinery and glycoprotein genes, we first mapped all human protein sequences (23,237) on the consensus phylogeny consisting of 29 phylogenetic levels (phylostrata - ps, Fig.1, Supplementary Data 1 and 2) (Domazet-Lošo and Tautz 2007, Domazet-Lošo and Tautz 2010a, Domazet-Lošo et al. 2017, Futo et al. 2021, Domazet-Lošo et al. 2024).

**Figure 1.**
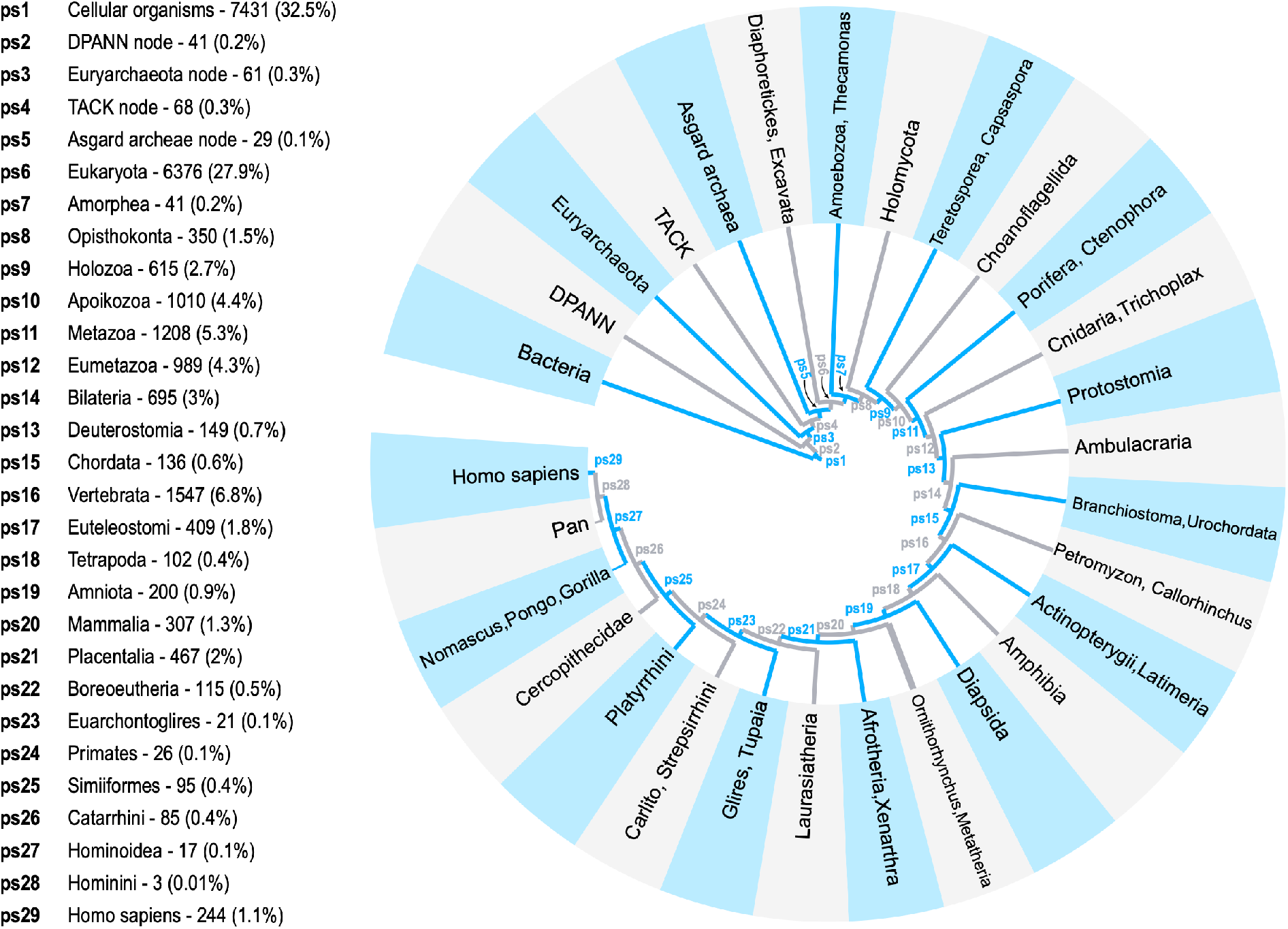
The consensus phylogeny used in the phylostratigraphic analysis. The condensed consensus tree covers divergence from the last common ancestor of cellular organisms to *Homo sapiens* as a focal organism. For a fully resolved tree see Supplementary Data 1. The tree is constructed by considering the importance of evolutionary transitions and availability of reference genomes. The internodes (29 phylostrata) that lead from the root of the tree to the focal species (*H. sapiens*) are marked by ps1-ps29. The numbers of *H. sapiens* genes traced to each phylostratum and corresponding percentages (in parentheses) are given after phylostrata names. We mapped in total 23,237 *H. sapiens* genes.

To get an overview of macroevolutionary patterns related to glycobiology we made a compilation of 673 human glycosylation machinery genes (see Methods). The majority of these genes (56%) map to the origin of all cellular orsganisms (ps1, Fig. 2), which is significantly more than expected by chance (odds ratio = 2.71, p = 1.29 x 10^-37^). The origin of cellular organisms (ps1, Fig. 2) is the only evolutionary period in which we detected the enrichment of glycosylation machinery genes indicating that the glycosylation is an ancient process common to all life. However, we detected also a substantial number of glycosylation machinery genes at the origin of eukaryotes (24%; ps6, Fig. 2) suggesting that in this evolutionary transition glycosylation pathways were further elaborated. Another 15% of glycosylation machinery genes we traced back to the period between the origin of Amorphea and Bilateria (ps7- ps13, Fig. 2). These numbers reveal that the origin of animal multicellularity as well as the body plan assembly of eumetazoans and bilaterians required new additions to glycosylation pathways. Notably, after the origin of amniotes (ps19, Fig. 2) no new glycosylation machinery genes were mapped on our phylostratigraphic map. Taken together, our phylostratigraphic profile of human glycosylation machinery genes suggests that glycosylation is an evolutionarily ancient process, fully established before the radiation of mammals (ps20, Fig. 2).

**Figure 2.**
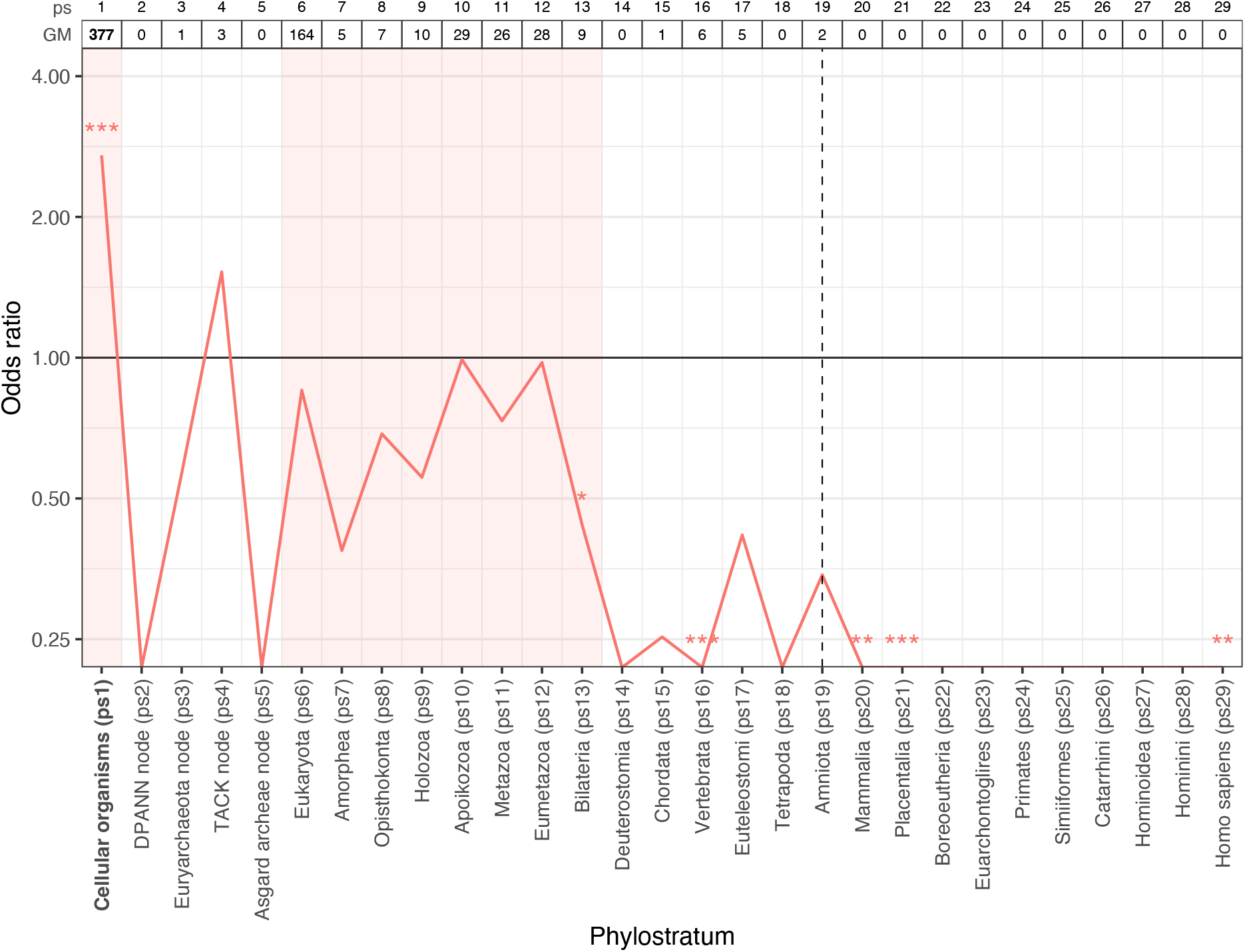
Phylostratigraphic analysis of *H. sapiens* glycosylation machinery genes. The x-axis shows 29 phylogenetic levels (phylostrata) of the consensus phylogeny with *H. sapiens* as a focal species (Fig.1). The table at top shows distributions of 673 human glycosylation machinery (GM) genes along phylostrata. The numbers in bold mark phylostrata with a substantial number of glycosylation machinery genes. The odds ratio chart shows deviation from the expected frequency. We tested the significance of glycosylation machinery genes enrichments and depletions using two-tailed hypergeometric test (\**p* < 0.05, \*\**p* < 0.01, \*\*\**p* < 0.001). Glycosylation machinery genes are only significantly enriched in ps1. However, the evolutionary origin of most glycosylation machinery genes could be traced to Cellular organisms (ps1), Eukaryota (ps6), and the period between Amorphea and Bilateria (ps7-ps13). These periods are shaded in red. The dashed vertical line marks the origin of amniotes (ps19). After this phylostratum no new glycosylation machinery genes emerged (ps20-ps29).

The reported results are based on the blastp e-value cutoff of 10^-3^ which has repeatedly been shown to be optimal in phylostratigraphic analysis (Domazet-Lošo and Tautz 2003; Domazet-Lošo et al. 2017; Moyers and Zhang 2018; Vakirlis et al. 2020, Futo et al. 2021, Domazet-Lošo et al. 2024). However, to test the stability of the observed phylostratigraphic patterns we recently introduced a test, where the analysis is repeated for a broad range of e-value cutoffs; e.g., between 1 and 10^-20^ (Futo et al. 2021). This sliding e-value protocol intentionally inflates false positive (e-values closer to 1) and false negative rates (e-values closer to 10^-20^) and thus tests the robustness of the initially observed phylostratigraphic pattern at 10^-3^ e-value cutoff (Futo et al. 2021). Our sliding e-value analysis confirmed the stability of the signal, which we initially found in ps1, in the full range of tested cutoff values (Supplementary Fig. S1). This result reassured us that the observed pattern at the 10⁻³ e-value reflects a genuine evolutionary imprint, rather than noise from the error rates of the blastp algorithm.

To get a more detailed insight into the evolutionary patterns of glycosylation machinery genes, we divided them into seven subgroups, which reflect their specific roles in glycobiology (Boutet et al. 2016, Reily et al. 2019). We then analyzed these subgroups independently using phylostratigraphic procedure (Fig. 3). The monosaccharide metabolism genes (MM) that contribute to glycosylation process are prevailingly mapped to the evolutionary oldest phylostrata (ps1) where they contribute to a strong enrichment signal (Fig. 3). This pattern suggests that the basic sugar metabolism necessary for glycosylation (i.e., monosaccharide activation) was present at the origin of cellular organisms. Comparably, genes that contribute to the N-glycosylation (NG) and O-glycosylation (OG) of proteins show a bimodal pattern with significant enrichments at the origin of cellular organisms (ps1, Fig. 3) and eukaryotes (ps6, Fig. 3). This enrichment profile suggests that protein glycosylation was present at the origin of cellular organisms (ps1), but also points to further innovations linked to these pathways at the origin of eukaryotes (ps6).

**Figure 3.**
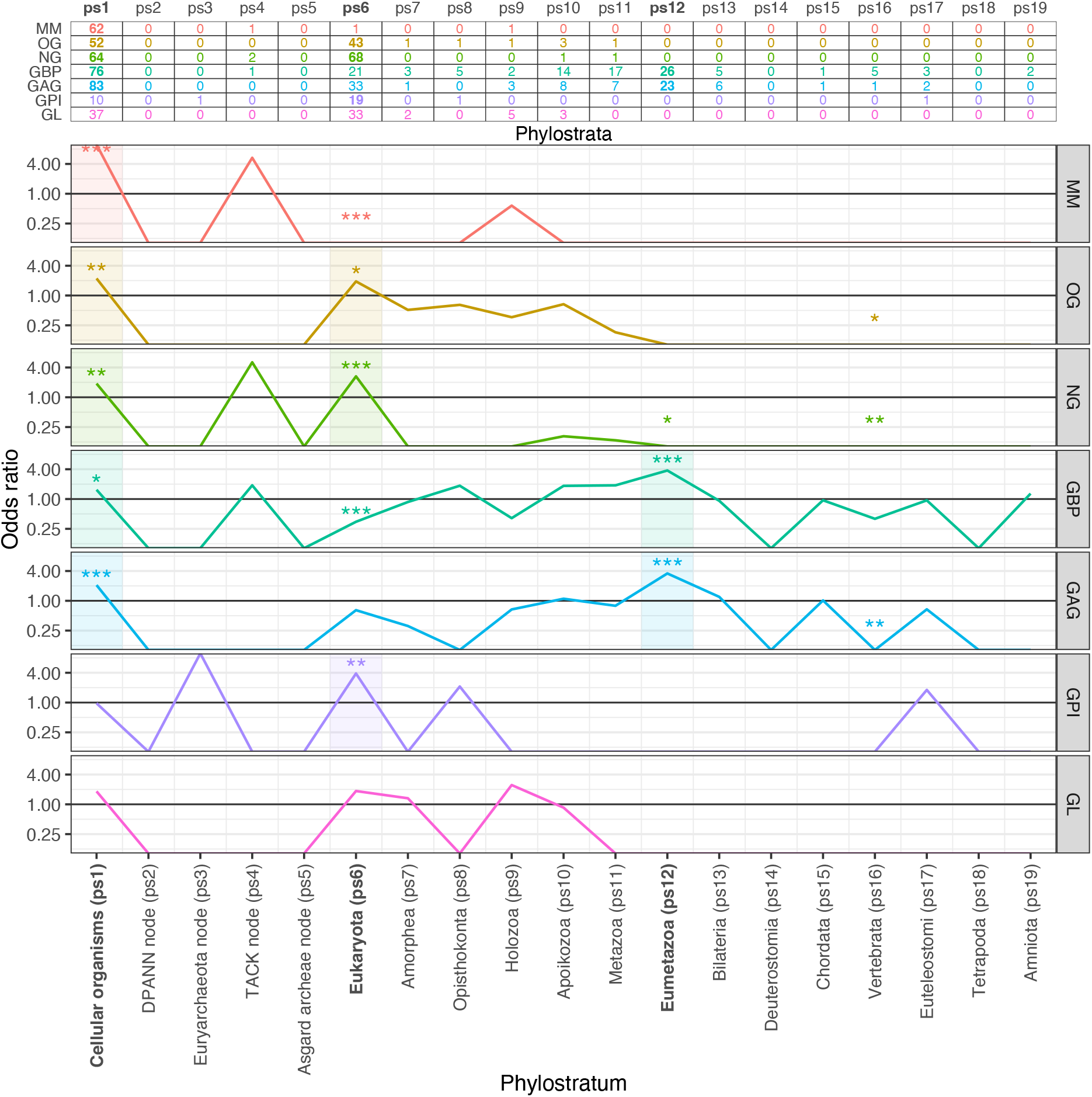
Phylostratigraphic maps of glycosylation machinery genes per functional groups. The table at top shows the distributions along 19 phylostrata of *H. sapiens* glycosylation machinery genes divided into seven specific glycosylation roles (monosaccharide metabolism - MM, O-glycosylation - OG, N-glycosylation - NG, glycan binding proteins - GBP, glycosaminoglycans - GAG, glycosylphosphatidylinositol anchors - GPI and glycolipids - GL). In contrast to Fig. 2, we here depicted only ps1 to ps19 because there are no new glycosylation machinery genes after the origin of amniotes (ps19, Fig. 2). The odds ratio charts show deviation from the expected frequency for every subgroup independently. We tested significance of the enrichments and depletions using two-tailed hypergeometric test (\**p* < 0.05, \*\**p* < 0.01, \*\*\**p* < 0.001). The shaded areas mark phylostrata with significant enrichments.

Genes involved in the synthesis of glycosaminoglycans (GAG) and glycan-binding proteins (GBP) produced very similar phylostratigraphic patterns (Fig. 3). Both groups of glycosylation machinery genes showed enrichment signals at the origin of cellular organisms (ps1, Fig. 3) and at the origin of eumetazoans (ps12, Fig. 3). The first signal at the origin of cellular organisms (ps1) suggests that the synthesis of glycosaminoglycans and the production of glycan-binding proteins have deep roots in evolutionary history. However, the second signal at the origin of eumetazoans (ps12) points that novel glycosaminoglycans and glycan-binding proteins played an important role in the emergence of the first metazoans. This is not surprising because glycosaminoglycans are the main component of the extracellular matrix and endothelial glycocalyx layer — the crucial elements of true tissues that determine their physical characteristics (Zeng et al. 2012; Lindahl et al. 2017). Similarly, glycan-binding proteins take an important role in cell adhesion, cell-cell interactions, cell-matrix interactions, and immune processes via self-nonself recognition; all of which are important functions of animal lifestyle (Taylor et al. 2015).

The glycosylphosphatidylinositol (GPI) anchor is a glycan-based posttranslational modification that allows modified proteins to be attached to the outer surface of the plasma membrane (Paulick and Bertozzi 2008, Vogt et a 2020, Beihammer et al. 2020). The glycosylation machinery involved in the formation of GPI anchored proteins showed a strong enrichment signal at the origin of eukaryotes (ps6, Fig. 3). This signal, in line with the observation that GPI modifications are found only in eukaryotes (Nosjean 1998, Kinoshita 2016), suggests that GPI anchor is a eukaryogenesis-linked innovation that facilitates protein trafficking within highly compartmentalized eukaryotic cells. In contrast, glycosylation machinery genes involved in glycolipid (GL) formation did not show any statistically significant enrichment (Fig. 3). Regardless of this, the vast majority of GL genes, similar to the other subgroups of glycosylation machinery genes, map to the origin of cellular organisms (ps1) or eukaryotes (ps6, Fig. 3). Finally, all enrichment profiles that we recovered from the subgroups of glycosylation machinery genes showed stability in sliding e-value analyses (Supplementary Fig. S2).

To compare evolutionary patterns of glycosylation machinery genes with their target proteins we mapped 4,565 human genes coding for glycoproteins (GP) onto the consensus phylogeny (Fig. 4). Around 38% of glycoproteins could be traced back to the ancestor of cellular organisms (ps1), which is significantly more than expected by chance (odds ratio = 1.30, p < 0.001). Apart from that, we detected significant enrichment signals in the evolutionary period that spans the origin of animals (Apoikozoa, ps10, Fig. 4) and bony vertebrates (Euteleostomi, ps17, Fig. 4). Approximately 37% of glycoproteins have their origin in this period with statistically significant odds ratio in the range between 1.26 and 2.32. These results corroborate the notion that glycosylation is an ancient process common to all life, because we found the overlapping enrichment of both types of genes, i.e., glycosylation machinery genes as well as glycoprotein genes, at the origin of cellular organisms (ps1, Fig. 2,4). In addition, the strong enrichment signals of glycoproteins in the span between the unicellular ancestors of animals (ps10, Fig. 4) to bony vertebrates (ps17, Fig. 4) suggests that the increased use of glycoproteins played an important role in the origin of the first animals and in their further radiation along the Cambrian and Ordovician geological periods (Schröder and Bosch 2016), which is likely connected with the immense energetic shift at the origin of animals (Kasalo et al. 2024). The sliding e-value analyses confirmed the robustness of these results (Supplementary Fig. S3).

**Figure 4.**
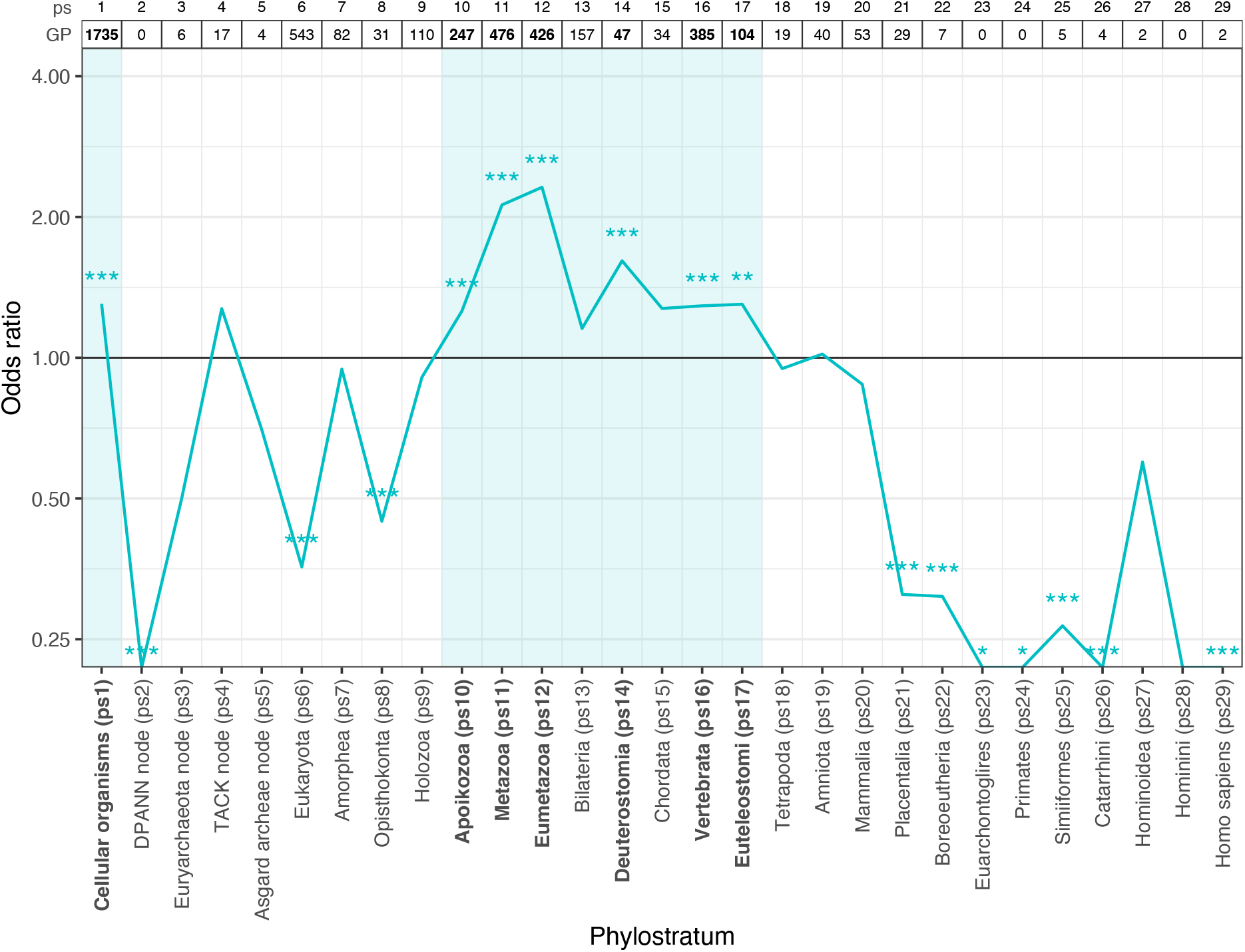
Phylostratigraphic analysis of *H. sapiens* genes coding for glycosylated proteins. The lower panel shows consensus phylogeny with *H. sapiens* as a focal species that contains 29 phylogenetic levels (phylostrata). We mapped all *H. sapiens* proteins (23,237) on the phylogeny (see Methods). The table at top shows distributions of genes which code for glycosylated proteins (GPs) and all genes along phylostrata. The odds ratio chart in the upper panel shows deviation from the expected frequency. We tested significance of the enrichments and depletions using two-tailed hypergeometric test (\**p* < 0.05, \*\**p* < 0.01, \*\*\**p* < 0.001). The evolutionary origin of most GP genes could be traced to ps1 (Cellular organisms) and ps6 (Eukaryota). GP genes are strongly enriched in ps1 (Cellular organisms) and in the evolutionary period that spans the origin of animals and bony vertebrates (Apoikozoa, ps10 to Euteleostomi, ps17). The gray shaded areas mark phylostrata with significant enrichments.

The vast majority of human glycosylated proteins (97%) in our dataset carry N-linked glycans. This sharply contrasts O-glycosylations which are much less frequent (7%) post-translational modification. By assuming that these large differences reflect the relative biological importance of these two post-translational modification types, and by considering that N-glycosylation is the best-studied protein glycosylation process, we focused our further analysis on N-glycosylation machinery. We first depicted the part of the N-glycosylation pathway that acts in the endoplasmic reticulum (ER) (Fig. 3). This subset includes 40 proteins that act directly in the biosynthetic pathway on the cytoplasmatic and luminal side of the ER membrane as well as those that are indirectly involved by providing activated monosaccharide blocks or are acting as membrane transporters (Fig. 5A). The N-glycosylation pathway starts with dolichol-phosphate biosynthesis or recovery at the cytosolic side of ER (Fig. 5A). We traced the evolutionary origin of three genes involved in these processes (DOLPP1, SRD5A4, and DOLK) to Eukarya (ps6) which suggests that they appeared for the first time in the last common ancestor of all eukaryotes (LECA); in line with the results of a previous study (Lombard 2016a).

**Figure 5.**
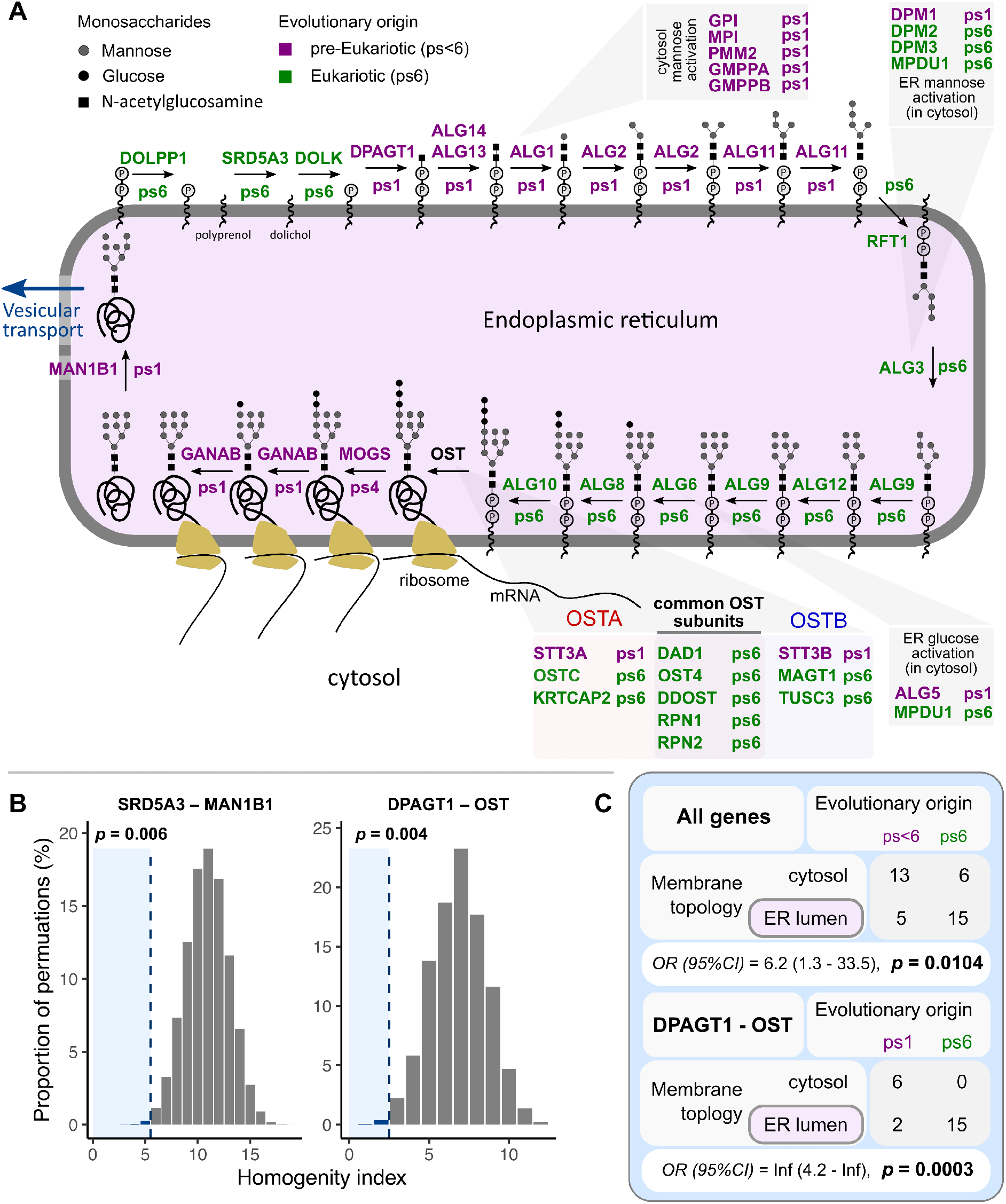
Biased evolutionary origin of N-glycosylation machinery genes in the endoplasmic reticulum. (**A**) The part of the human N-glycosylation biosynthetic pathway which is linked to the endoplasmic reticulum (ER). The N-glycosylation machinery proteins (40 genes) are represented with HGNC gene symbols, which are followed by corresponding phylostrata of their evolutionary origin (ps). Genes (proteins) that emerged at origin of Cellular organisms (ps1), plus MOGS gene that we traced to TACK Archeae (ps4), are in violet. Genes (proteins) that emerged at the origin of Eukaryotes (ps6) are in green. (**B**) Empirical probability distribution of homogeneity indices (HI) calculated by positional permutation of phylostrata. Dashed line represents the observed HI value, while *p*-value is probability of obtaining exactly that or a smaller HI value. (**C**) Contingency tables show the distribution of genes across two factors: evolutionary origin and membrane topology. We tested nonrandom associations between evolutionary origin and membrane topology using Fisher’s exact test. Odds-ratio (OR) values and 95% confidence (CI) intervals are shown.

The next step in the N-glycosylation process is the addition of the first glycan on the dolichol-phosphate carrier catalyzed by a glycosyltransferase (DPAGT1) on the cytosolic side of the ER membrane (Fig. 5A). After this modification, a series of glycosyltransferases (ALG13, ALG14, ALG1, ALG2, and ALG11) sequentially add monosaccharides to yield the Man5GlcNAc2 structure. All these genes are mapped to ps1, suggesting their evolutionary occurrence in the last common ancestor of all cellular organisms (Fig. 5A). This newly synthesized glycan structure, which is attached to the dolichol carrier, is then flipped by the flippase (RFT1) into the lumen of the ER (Fig. 3A). The N-glycosylation process proceeds further on the luminal side of the ER membrane where a series of glycosyltransferases (ALG3, ALG9, ALG12, ALG6, ALG8, and ALG10 genes) add additional glycans to reach the final Glc3Man9GlcNAc2 structure common to all eukaryotes. All these proteins that act on the luminal side of the ER membrane are mapped to the origin of Eukaryota (ps6) (Fig. 5A).

In the next step, the oligosaccharyltransferase (OST) protein complex transfers the glycan from the dolichol carrier to the polypeptide (Fig. 5A). This protein complex is assembled from five constant and three variable subunits. We traced all these subunits to the origin of Eukaryota (ps6), with the exception of core catalytic subunit SST3, which we mapped at the origin of cellular organisms (ps1). This phylostratigraphic pattern suggests that the OST complex evolved during eukaryogenesis by the addition of new regulative subunits to the ancient core catalytic protein (SST3). Once an oligosaccharide is attached to a protein, the processing of the glycan begins with the removal of the glucose residues with two glucosidases (MOGS and GANAB). We traced MOGS to the origin of TACK/Asgard archaea (ps4) and GANAB to the origin of cellular organisms (ps1). When the glycoprotein is finally properly folded, α- mannosidase I (MAN1B1), which we traced to cellular organisms (ps1, Fig. 5A), removes one mannose from the central glycan arm which signals that the glycoprotein is ready for transport outside the ER, usually to the Golgi apparatus (Moremen et al. 2012).

By looking at the phylogenetic distribution of N-glycosylation ER proteins (Fig. 5A), we found that essentially all of them come from only two evolutionary periods: cellular organisms (ps1) or Eukaryotes (ps6). Besides this binary evolutionary origin of N-glycosylation ER proteins, which follows the global evolutionary pattern of N-glycosylation genes (Fig. 3), we noticed that the proteins of the same evolutionary origin tend to have adjacent positions along the biosynthetic pathway (Fig. 5A). For instance, there is a block of seven consecutive biochemical steps (DPAGT1 to ALG11) where all proteins have evolutionary roots at the origin of cellular organisms (ps1) (Fig. 5A). This block is followed by an evolutionary younger one (RFT1- ALG10) that contains eight biochemical steps where all proteins have phylogenetic roots at the origin of eukaryotes (ps6) (Fig. 5A).

To assess this phenomenon quantitatively, we devised a positional homogeneity index (HI) which estimates the clustering strength of identical phylostrata along a biosynthetic pathway (see Methods). Low HI values reflect an extensive positional grouping of the identical phylostrata, whereas high HI values point to a scattered arrangement of phylostrata along the biosynthetic pathway. By applying this measure to the sequence of all N-glycosylation reactions in the ER, we found a very low HI value that signals the biased grouping of phylostrata (HI = 5, Fig. 5B). To test if this biased grouping is statistically significant, we compared the observed HI value to the distribution of all possible HI values in the ER part of the N-glycosylation pathway (see Methods). We found that is extremely unlikely that such strong positional clustering of identical phylostrata occurred by chance (p = 6.97×10^-4^, Fig. 3B). We obtained similar results if we focused only on the core segment of the ER N-glycosylation pathway from DPAGT1 to OST, which starts with the first attachment of a monosaccharide to the lipid carrier and ends with the transfer of the glycan to the polypeptide chain. (HI = 2, p= 6.99×10^-4^, Figure 5B).

Besides these longitudinal clustering biases along the ER N-glycosylation pathway, we noticed that phylostrata were not randomly distributed with respect to the position of N-glycosylation machinery on the two sides of the ER membrane. The reactions that occur at the cytosol side of the ER membrane tend to use N-glycosylation machinery proteins that we traced to the origin of cellular organisms (ps1) (Fig. 5A, C). Conversely, the reactions that unfold at the luminal side of the ER membrane preferentially rely on the N-glycosylation machinery proteins that we traced to the origin of eukaryotes (ps6) (Figure 5A, C). This biased arrangement is significant when all genes are analyzed (Fisher’s exact test, *p* = 2.7×10^-3^) as well as when only the core part of the N-glycosylation pathway from DPAGT1 to OST is considered (Fisher’s exact test, *p* = 2.08×10^-4^). These results are quite stable in a broad range of e-value cutoffs (Supplementary Fig. S4).

After reaching Golgi, N-glycans go through further processing that involves coordinated action of glycosidases and glycosyltransferases that gives rise to three main classes of glycans: oligomannosidic, hybrid, and complex glycans. However, some glycans can completely skip these modifications (Frappaolo et al. 2020). The Golgi apparatus is organized into discrete cisternae, each containing a distinct subset of N-glycosylation machinery proteins which include glycosidases, glycosyltransferases, and nucleotide sugar transporters that provide glycosyltransferases with nucleotide sugars as glycosyl donors (Frappaolo et al. 2020, Mikkola et al. 2020, Hadley et al. 2019) (Figure 6A). Similar to the ER, we found that N-glycosylation proteins in Golgi are also assigned only to two phylostrata: cellular organisms (ps1) and eukaryotes (ps6) (Figure 6A).

**Figure 6.**
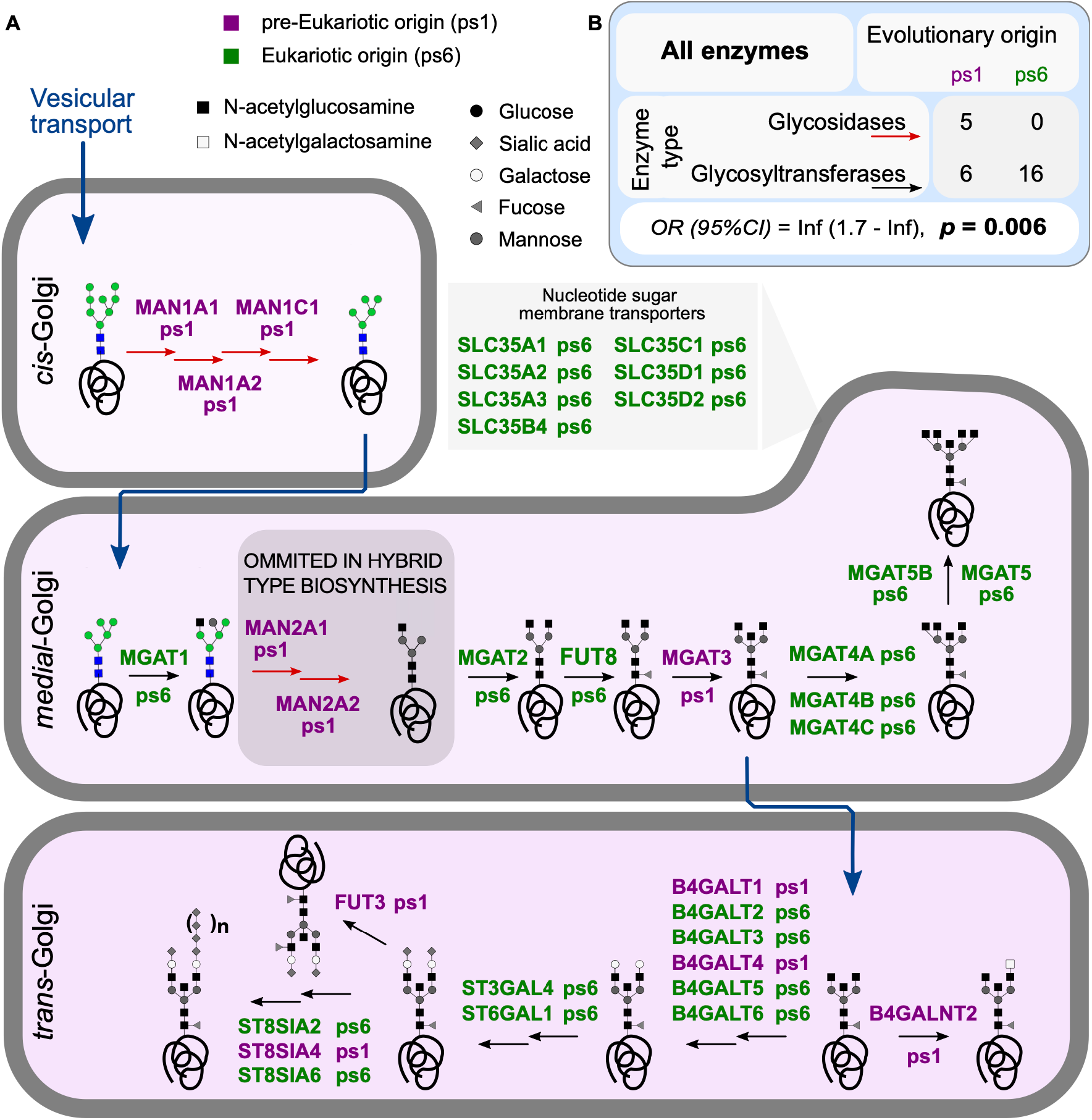
Biased evolutionary origin of N-glycosylation enzyme types in the Golgi apparatus. (**A**) The part of the human N-glycosylation biosynthetic pathway which is linked to the Golgi apparatus. N-glycosylation machinery proteins (27) are represented with HGNC gene symbols, which are followed by corresponding phylostrata of their evolutionary origin (ps). Genes (proteins) that emerged at origin of Cellular organisms (ps1) are in violet. Genes (proteins) that emerged at the origin of Eukaryotes (ps6) are in green. (**B**) The contingency table shows the distribution of genes across two factors: evolutionary origin and enzyme type. We tested nonrandom associations between evolutionary origin and enzyme type using Fisher’s exact test. The odds-ratio (OR), *p*-value and 95% confidence (CI) intervals are shown.

We traced seven nucleotide sugar membrane transporters acting in Golgi (SLC35 gene family) to the origin of eukaryotes (ps6) (Figure 6A). In contrast, we tracked all five glycosidases (mannose trimming enzymes), located in the cis (MAN1A1, MAN1A2, and MAN1C1) and medial-Golgi (MAN2A1, and MAN2A2), to the origin of cellular organisms (ps1) (Figure 6). On the other hand, 22 glycosyltransferases have binary phylogenetic origin. The majority of them (16) could be traced to eukaryotes (ps6) and the rest (6) to the origin of cellular organisms (ps1) (Figure 6). In contrast to the ER, the Golgi N-glycosylation pathway is not linear and does not have membrane polarity as all glycosidases or glycosyltransferases reactions occur in the Golgi lumen. These properties prevented us from applying heterogeneity index or testing location biases similar to those applied in the ER (Figure 5). However, we noticed an obvious bias in the phylogenetic origin of proteins acting in Golgi depending on their functional roles, where all glycosidases could be traced back to the origin of cellular organisms (ps1) and the majority of glycosyltransferases to the origin of eukaryotes (ps6). There is a low probability to observe such distribution by chance, (Fisher’s exact test, p = 5.72×10^-3^) (Figure 4B), and the pattern is stable for e-values above 10^-3^ (Supplementary Fig. S5).

## Discussion

Our global phylostratigraphic analysis of glycosylation machinery genes in humans revealed that glycosylation is an evolutionary old process most likely common to all life. However, glycosylation machinery pathways relevant to humans were established along protracted macroevolutionary time via three important periods. The first one is the origin of cellular organisms (ps1, Fig. 2,3) where we traced most of the human glycosylation machinery. The second is the origin of eukaryotes (ps6, Fig. 2,3) and the third one covers eukaryotic radiation and early metazoan diversification (ps7-ps13, Fig. 2,3). Finally, the complete glycosylation machinery, on which extant humans depend, was most likely fully established before the origin of mammals (ps20, Fig. 2,3). However, it is interesting that we traced the comparably small number of glycosylation machinery genes to the archaea lineage (ps2-ps5, Fig. 2,3). This pattern suggests that glycosylation in humans predominantly relies on the glycosylation machinery that was already present at the origin of cellular organisms (ps1) and not on some archaea-specific innovations (ps2-5). However, this sharply contrasts the subsequent evolutionary period, the onset of eukaryotes, where we detected the burst of new glycosylation machinery genes (ps6, Fig. 2,3). This pattern creates a polar distribution in the analysis of the N-glycosylation pathway where glycosylation machinery genes are either common to all cellular life (ps1) or specific to eukaryotes (ps6) (Fig. 5,6).

This binary evolutionary origin of N-glycosylation machinery genes (ps1 vs. ps6) is a useful marker which allowed us to look for potential biases in the distribution of N-glycosylation machinery genes on the endoplasmic membrane. Surprisingly, we found a non-random distribution where N-glycosylation genes specific for cellular organisms (ps1) are predominantly located on the cytosolic side and those specific for eukaryotes (ps6) on the luminal side of the ER membrane (Fig. 5). This biased distribution has important evolutionary implications because it suggests that the lumen of the endoplasmic reticulum is a eukaryotic innovation which was devised by the invagination of the prokaryotic plasma membrane (Fig. 7). Bacterial and archaeal N-glycosylation machinery is always positioned at the innermost cell membrane where glycan synthesis occurs on the cytoplasmic side of the membrane. When completed, glycan is flipped to the extracellular or periplasmatic space (Fig. 7A). If we assume that this cross-membrane directionality of N-glycosylation is preserved in eukaryotes, then the most likely explanation for the fact that in eukaryotes the flipping of the Man5GlcNAc2 structure occurs towards the ER lumen is that ER emerged by the invagination of the prokaryotic innermost membrane that contained N-glycosylation machinery (Fig. 7). Our finding that N-glycosylation genes located on the cytoplasmic side of the ER membrane tend to be common to all cellular organisms (ps1), and that those at the luminal side are preferentially of eukaryotic origin (ps6), further corroborate this view (Fig. 5).

**Figure 7.**
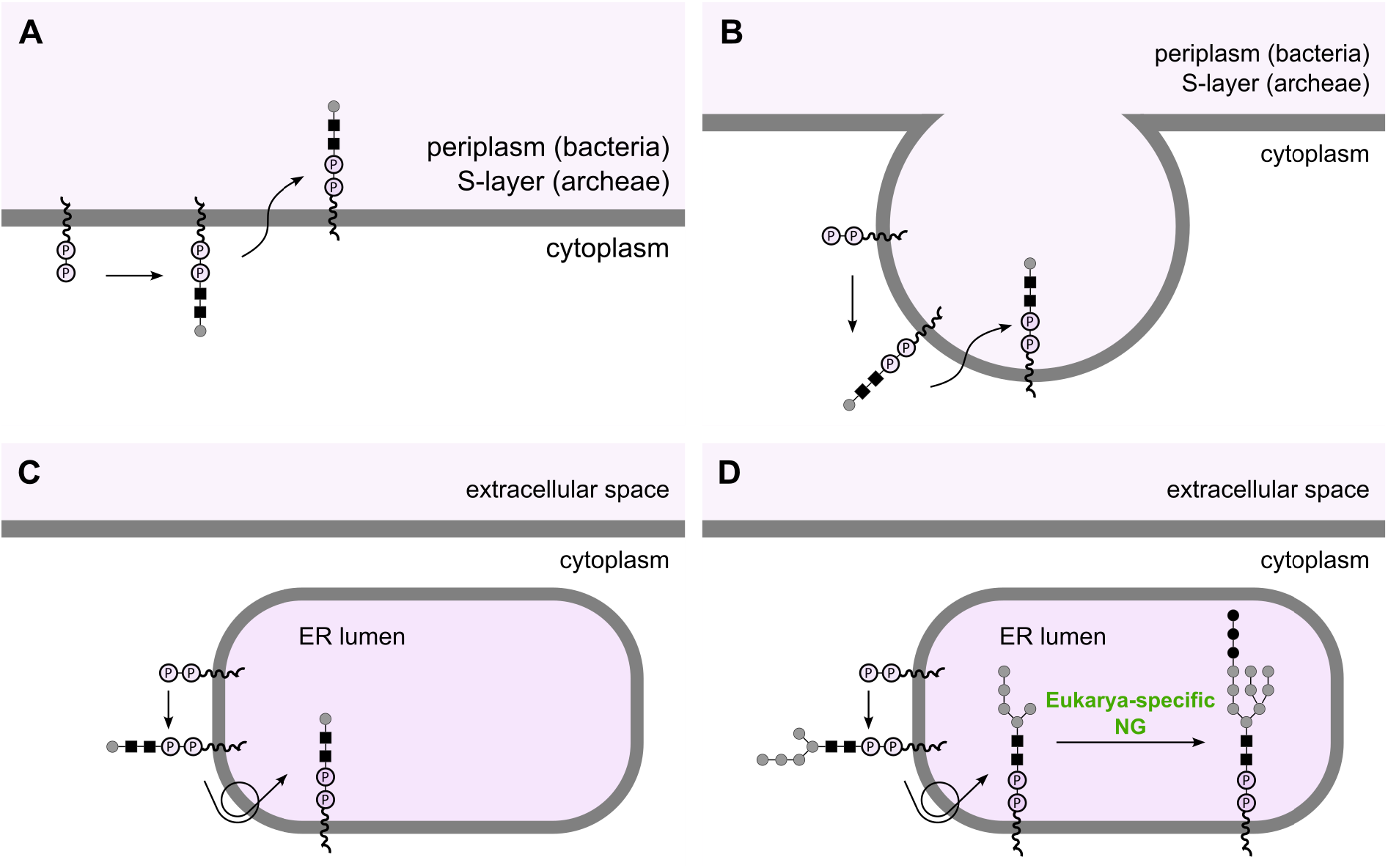
A plausible evolutionary scenario for the placement of N-glycosylation machinery in the endoplasmic reticulum. **(A)** Schematic representation of the N-glycosylation pathway in bacteria and archaea. N-glycosylation occurs on the cytoplasmatic side of the innermost cell membrane. After the glycan synthesis is completed on the membrane-bound lipid carrier the resulting glycan is enzymatically flipped to the periplasm or S-layer of bacteria and archaea. **(B)** The plausible invagination of the prokaryotic innermost membrane that contained N-glycosylation machinery. **(C)** If this forming vesicle pinched off the cell membrane, then the residing N-glycosylation machinery gains the spatial position identical to that present in the eukaryotic ER. **(D)** During eukaryogenesis new genes were added to the N-glycosylation pathway which further process glycans inside the lumen of ER.

However, the question arises regarding the capability of extant and ancestral prokaryotes to form intracellular membrane vesicles (IMVs). In contrast to extracellular membrane vesicles (EMVs) which have been recently extensively studied in prokaryotes (Toyofuku et al. 2019, Gill et al. 2019), IMVs are a much less explored feature. Nevertheless, it is clear that bacterial cells could also form IMVs (Lonhienne et al. 2010, Forterre and Gribaldo 2010, Saier 2014, Grant 2018, Kapteijn et al. 2022, Flood et al. 2022). For instance, it was shown that cell wall-deficient bacteria can encapsulate extracellular material by invagination of the cytoplasmic membrane by an endocytosis-like process (Kapteijn et al. 2022). This is an especially suggesting finding because wall-deficient bacteria are considered a model of early cellular life before the invention of the cell wall (Forterre and Gribaldo 2010, Errington et al. 2016, Kapteijn et al. 2022).

Another important question relates to the evolutionary origin of the N-glycosylation genes that in our analysis map to the cellular organisms (ps1). In principle, there are two contexts when these genes are placed at ps1. In the first one, these genes could have a significant match to bacteria only, while in the second one, these genes could have significant matches to both bacteria and archaea. A situation where most of the N-glycosylation genes that are placed at ps1 have significant matches only to bacteria would indicate that the evolutionary oldest part of the eukaryotic N-glycosylation pathway stems from the bacterial ancestor. In turn, this would imply that bacteria and archaea evolved N-glycosylation independently. Conversely, a situation where most of the N-glycosylation genes that are placed at ps1 have significant matches to both bacteria and archaea would suggest that already LUCA possessed basic N-glycosylation machinery that was then inherited in all subsequent lineages.

To test for these possibilities, we created a heatmap of the best matches per phylostratum for N-glycosylation genes (Supplementary Fig. S6). This analysis revealed that most of N-glycosylation genes mapped to ps1, which act in ER, have significant matches in both bacteria and archaea, suggesting that the basic N-glycosylation machinery was already present in LUCA and that all diverging lineages were built on this ancient core. A notable exception is ALG1, MPI, and PMM2 which seem to have significant matches only in bacteria. Similarly, the majority of genes that map to ps1 in Golgi have significant matches exclusively in bacteria (Supplementary Fig. S6). Together, this gives some credence to the idea that eukaryotic N-glycosylation genes that map to ps1 were actually inherited from bacterial ancestor, in line with the notion that eukaryotic membranes are of bacterial origin (López-García and Moreira 2020). However, to fully resolve this issue it would be useful in the future to perform a detailed phylogenetic analysis of individual N-glycosylation ps1 genes.

Our heatmap analysis also reveals that N-glycosylation machinery is largely preserved in eukaryotic side-branches along human lineage (x-axis, ps6-ps29, Supplementary Fig. S6). However, there are also some exceptions. The most striking example is the loss of Golgi glycosyltransferase genes in fungi (x-axis, ps8, Supplementary Fig. S6). This finding agrees with the observed lack of galactose and N-acetylglucosamine residues in glycan structures found in fungi (Chung et al. 2017).

Taken together, our study showed that glycosylation in *Homo sapiens* is an ancient feature, with a biphasic origin that comprises the origin of cellular organisms and the origin of eukaryotes. Despite this ancient history of glycosylation machinery, the usage of protein glycosylation intensified with the development of complex multicellular organisms, probably linked to selective pressures related to self-nonself recognition and improved coordination between the differentiated cells. Finally, using N-glycosylation pathway as a marker, we provide some support for the idea that endoplasmic reticulum evolved through invagination of the prokaryotic cell membrane that possessed the N-glycosylation pathway.

## Methods

### Glycosylation machinery genes and glycoproteins

We compiled a list of glycosylation machinery (GM) genes from several sources: the supplementary table of Moremen et al. 2012, KEGG pathways related to glycosylation in *H. sapiens* (hsa00510, hsa00511, hsa00512, hsa00514, hsa00515, hsa00531, hsa00532, hsa00533, hsa00534, hsa00563, hsa00601, hsa00603, hsa00604) (Kanehisa and Goto 2000), *H. sapiens* entries in Carbohydrate Active enZYmes (CAZY) database (Cantarel et al. 2009), genes listed in Varki et al. 2015 and the supplementary table in Schjoldager et al. 2020. We also retrieved *H. sapiens* orthologues of mouse glycosylation genes listed in Moreman et al. (2012). The obtained dataset was manually reviewed for duplicates and pseudogenes which finally yielded a total of 673 glycosylation machinery genes with unique Ensemble gene IDs (Supplementary Data 3).

In addition, collected a list of glycoproteins (GP) from UniProtKB/Swiss-Prot database (Boutet et al. 2016 using the following query: “annotation:(type:carbohyd) AND reviewed:yes AND organism:"Homo sapiens (Human) [9606]"”. In this way, we obtained all the reviewed *H. sapiens* proteins that show some evidence to possess a posttranslational modification of glycosylation type. The final GP dataset included a total of 4565 genes with unique Ensemble gene IDs (Supplementary Data 3).

### Consensus phylogeny and reference genomes

We constructed a consensus tree of 503 organisms representing the diversity of major lineages that lead from the origin of cellular organisms to *Homo sapiens*, resulting in 29 phylogenetic levels (phylostrata) (Fig. 1, Supplementary Data 1 and 2). For phylogenetic relationships between taxa, we followed the relevant literature (Wang et al. 2009, Irisarri et al. 2017, Hughes et al. 2018, Regier et al. 2010, Misof et al. 2014, Shen et al. 2016, Berbee et al. 2017, Morris et al. 2018). Reference genomes, i.e., thier corresponding amino acid sequences, of all organisms included in the phylogeny were downloaded mostly from the Ensembl nad NCBI databases. Proteomes of choanoflagellates were retrieved from Richter et al. (2018). Prior to the phylostratigraphic analysis, the proteomes were processed to keep only the longest splicing variant of each gene. Other details on the construction of a phylostratigraphic database are described in Domazet-Lošo et al. (2024).

### Phylostratigraphic analysis

The theoretical background and procedure of phylostratigraphic analysis has been previously described (Domazet-Lošo and Tautz 2008; Tautz and Domazet-Lošo 2011; Domazet-Lošo et al. 2017; Domazet-Lošo et al. 2024). Briefly, the longest splicing variant of every *H. sapiens* gene was compared to the reference database using the blastp algorithm with the e-value cutoff equal to 10^-3^. The obtained sequence similarity search results, along with the consensus phylogeny, were used to estimate the evolutionary origin (phylostratum) of all human genes.

The frequency distributions of studied genes (GM and GP) were compared to the frequency distribution of all human genes across phylostrata (expected distribution). In all graphical representations the ratio of these frequencies is shown as log odds ratios. Deviations from the expected frequencies at each phylostratum were tested using a two-tailed hypergeometric test. The obtained p-values were corrected for multiple testing using the Benjamini-Hochberg method. To account for a potential sequence similarity search bias, we applied a sliding e-value protocol (Futo et al. 2021) where we repeated sequence similarity searches and all downstream calculations using a range of e-value cut-offs (1, 10^-1^, 10^-2^, 10^-5^, 10^-10^, 10^-15^, and 10^-20^).

## Statistical analyses

To test whether there is an association between the evolutionary origin of genes coding for enzymes of the N-glycosylation pathway acting on the endoplasmic reticulum and membrane topology, we generated a contingency table with these categories and tested observed biases using Fisher’s exact test. We used the same approach in the Golgi to test the association between the evolutionary origin and enzyme type.

To assess if there is an association between the order of enzymes of the N-glycosylation pathway in the ER and their evolutionary origin, we devised the homogeneity index (HI). For an ordered list of N genes with assigned evolutionary origin (phylostratum), HI is calculated by iterating from the 2nd to the N-th element of the list and comparing every element with the one immediately preceding it. If the evolutionary origin of the two elements is not equal, the index is increased by 1. In essence, a high HI means that the sequence of enzymes is disordered with respect to their evolutionary origin, while a sequence in which the enzymes of the same evolutionary origin are clustered together will result in a low HI.

To test whether the observed biased grouping of enzymes of the N-glycosylation pathway in the ER is statistically significant, we generated all possible permutations of evolutionary origins and calculated the HI for each permutation. We arranged the calculated HI values into bins containing identical values. We divided the number of elements in each bin with the total number of permutations, resulting in the empirical mass density function of HIs. Finally, we calculated the p-value by summing the probabilities of bins containing the HI values equal to or lower than the observed one. All statistical analyses were performed using Python and R (R Core Team 2021). Figures were created using R packages “ggplot2” and “cowplot” (Wickham 2016; Wilke 2020).

**Author Contributions:** G.K., G.L. and T.D.-L. initiated the study. D.K., G.K. and T.D.-L. conceptualized the analyses, D.K., N.Č., M.D.-L., and N.K. performed the analyses. D.K. prepared the figures and tables for publication. D.K., G.K. and T.D.-L. wrote the manuscript with contribution of all authors.

## Data availability

All data are available in the main text or the supplementary materials.

## Acknowledgments

We thank M. Futo, A. Tušar, S. Koska and D. Franjević for discussions. This work was supported by the Croatian Science Foundation under the project IP-2016-06- 5924 (T.D.-L.), the City of Zagreb (T.D.-L.), the Adris Foundation (T.D.-L.), the European Regional Development Fund KK.01.1.1.01.0009 DATACROSS (M.D.-L., T.D.-L.). We used the computational resources of the University Computing Center - SRCE (Padobran) and the Institute Ruđer Bošković.

## Competing interests

The authors declare no competing interests.

---

**Supplementary Figure S1.**
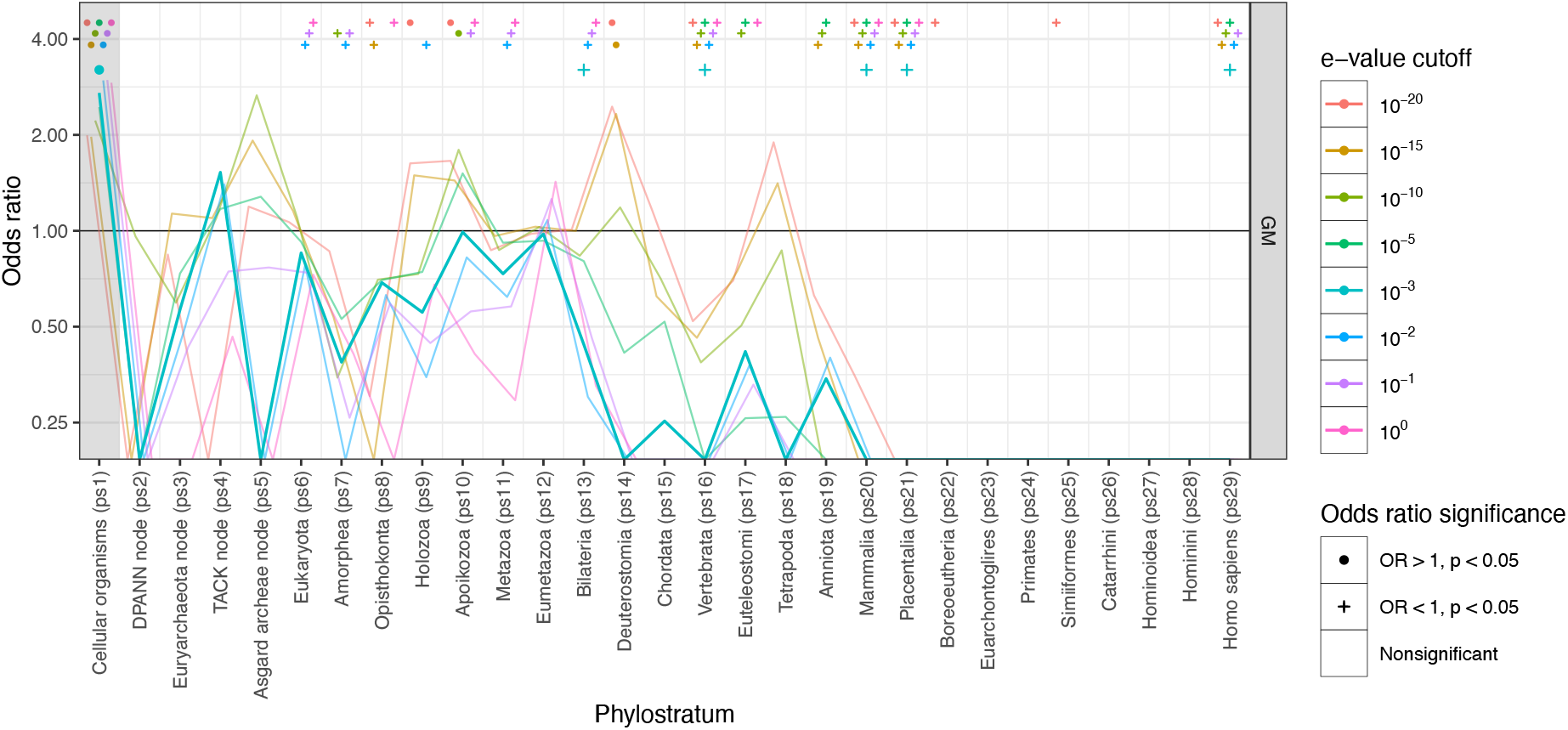
Testing the robustness of phylostratigraphic patterns by sliding e-value test (all glycosylation machinery genes). Odd ratio profiles were calculated for e-value cutoffs in the range between 1 and 10^-20^. Circles at the top of the panel mark significant enrichments, while plus signs mark significant depletions tested by a two-tailed hypergeometric test (*p* < 0.05). The profile shows a stable signal at ps1 (gray shaded area) where statistically significant gene enrichment was initially detected at the default 10^-3^ e-value cutoff (Fig. 2).

**Supplementary Figure S2.**
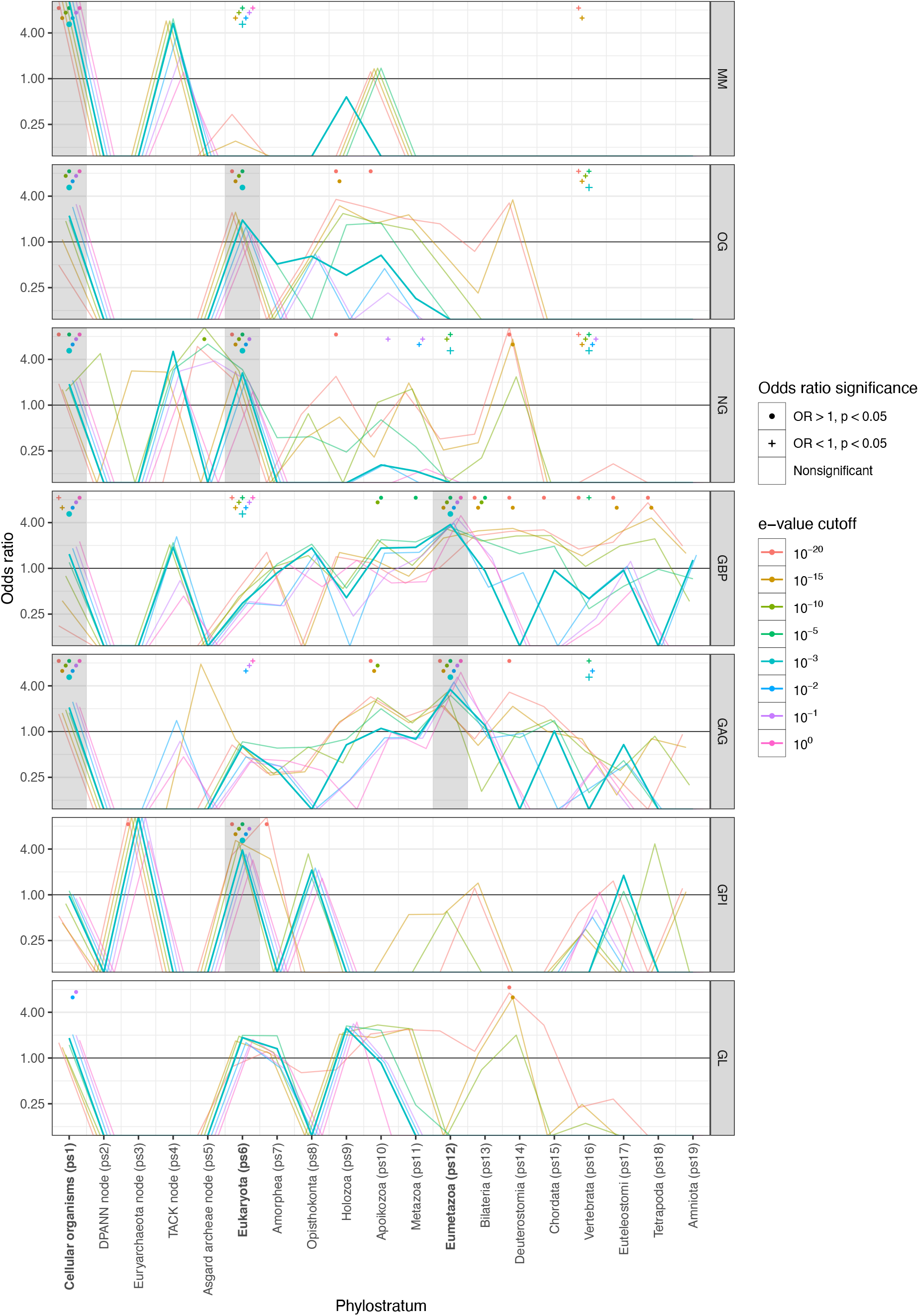
Testing the robustness of phylostratigraphic patterns by sliding e-value test (glycosylation machinery subgroups). The odds ratio charts along 19 phylostrata are shown for *H. sapiens* glycosylation machinery genes divided into seven specific glycosylation roles (monosaccharide metabolism - MM, O-glycosylation - OG, N-glycosylation - NG, glycan binding proteins - GBP, glycosaminoglycans - GAG, glycosylphosphatidylinositol anchors - GPI and glycolipids - GL). The odds ratio profiles were calculated for e-value cutoffs in the range between 1 and 10^-20^. Circles at the top of the panels mark significant enrichments, while plus signs mark significant depletions tested by two-tailed hypergeometric test (*p* < 0.05). Gray shaded areas highlight phylostrata where statistically significant gene enrichment signals were initially detected at the default e-value of 10^-3^ (Fig. 3). In all these phylostrata we find statistically significant gene enrichment signals in the full range of e-value cutoffs (from 1 to 10^-20^), with the exception of O-glycosylation signal at ps6 where enrichments were significant in the range between 10^-3^ and 10^-20^.

**Supplementary Figure S3.**
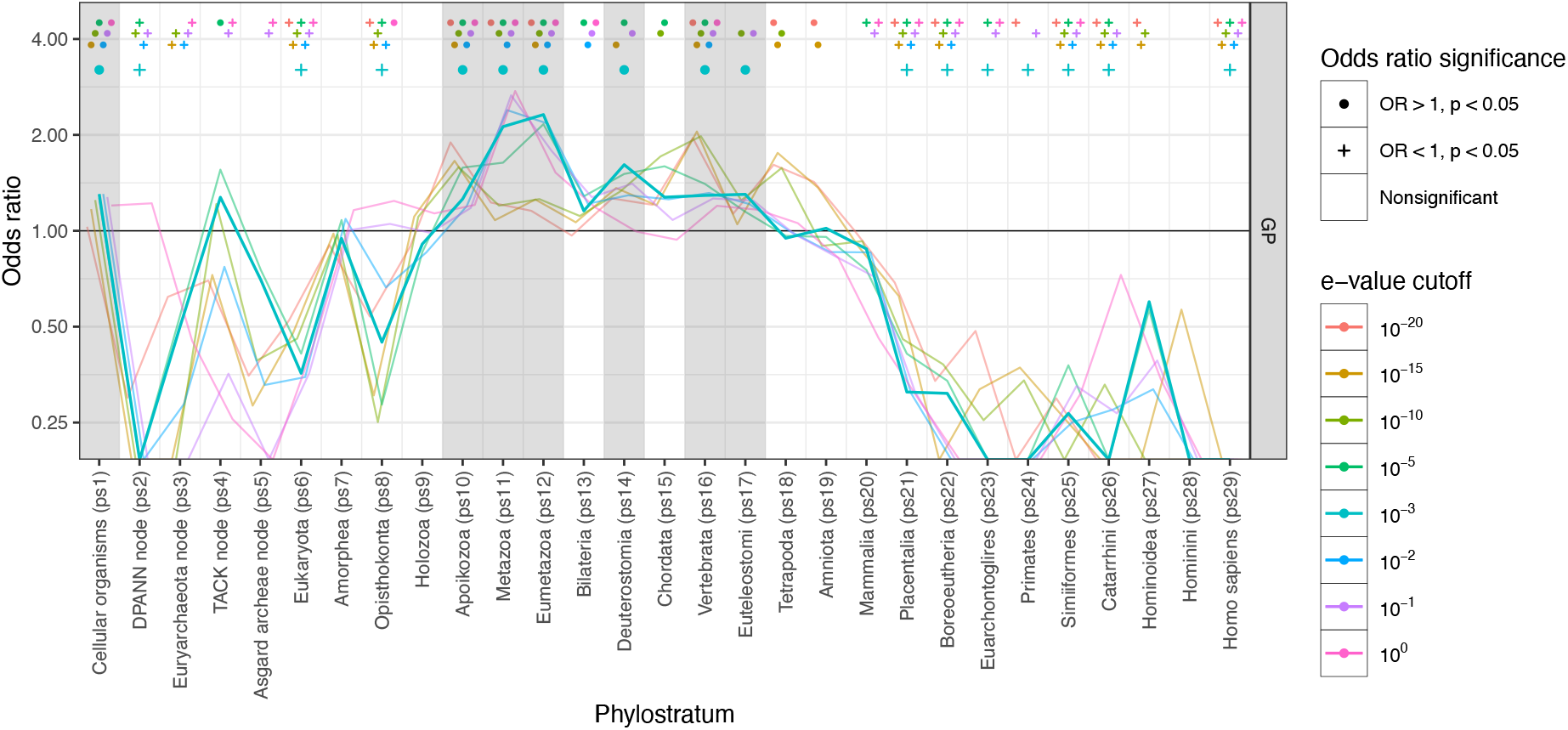
Testing the robustness of phylostratigraphic patterns by sliding e-value analysis (glycosylated proteins). Odd ratio profiles were calculated for e- value cutoffs in the range between 1 and 10^-20^. Circles at the top of the panels mark significant enrichments, while plus signs mark significant depletions tested by two-tailed hypergeometric test (*p* < 0.05). Gray shaded areas highlight phylostrata where statistically significant gene enrichment signals were initially detected at the default e-value of 10^-3^ (Fig. 3). In phylostrata Cellular organisms (ps1), Apoikozoa (ps10), Metazoa (ps11), Eumetazoa (ps12) and Vertebrata (ps16) we find statistically significant gene enrichment signals in the full range of e-value cutoffs (from 1 to 10^-20^), with the exception that the enrichments in ps1 at 10^-20^ and in ps11 at 10^-15^ were not significant.

**Supplementary Figure S4.**
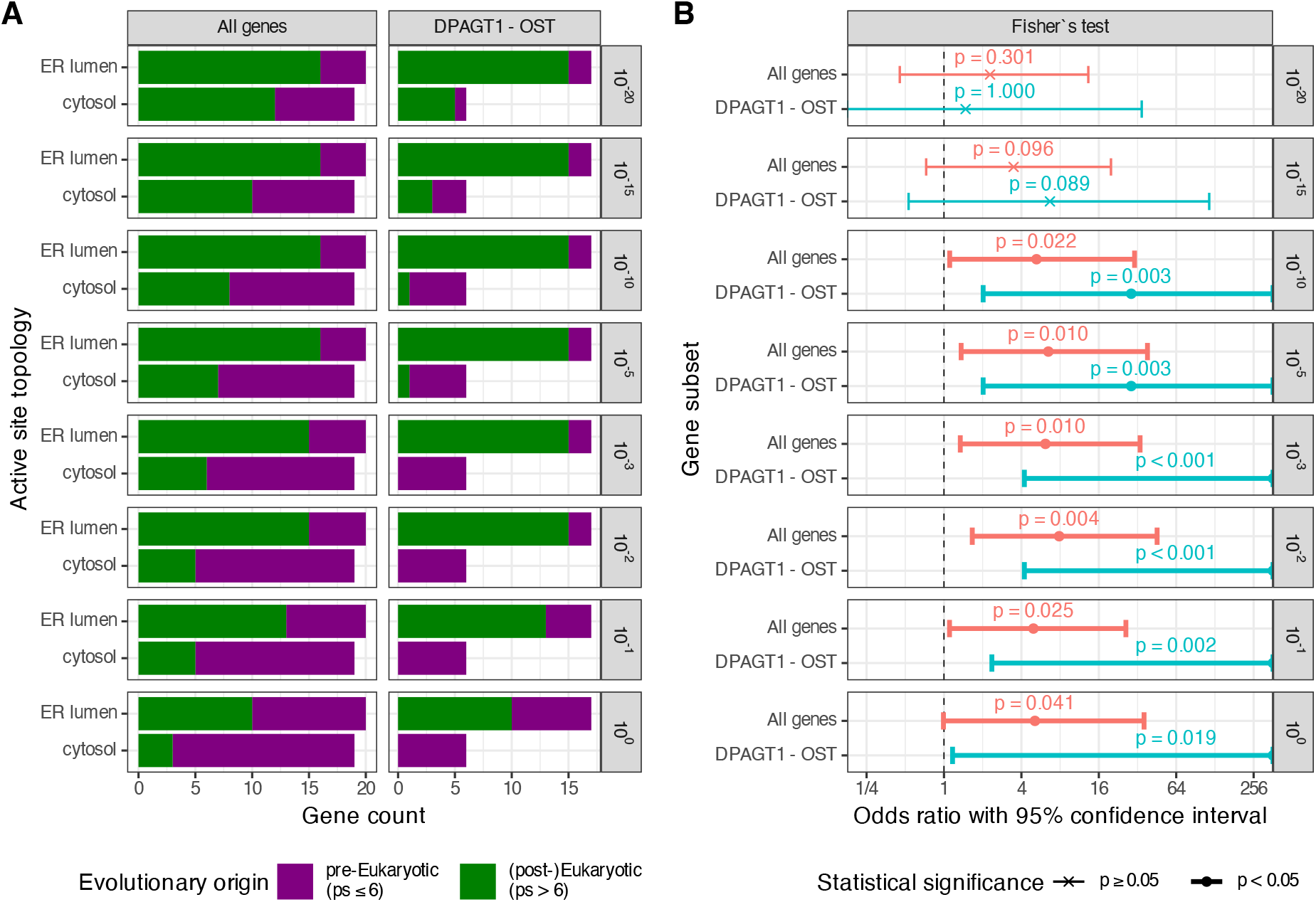
Testing the robustness of biases in the contingency tables by sliding e-value analysis (Figure 5C). We detected statistically significant bias in contingency tables that compare evolutionary origin and membrane topology of N-glycosylation genes in the endoplasmic reticulum (ER). (A) Proteins positioned on the cytosolic side of the membrane of the ER tend to be evolutionary older (Cellular organisms, ps1) in contrast to the proteins positioned on the luminal side of the membrane which tend to be evolutionary younger (Eukaryota, ps6). This bias was present in the full range of tested e-values (from 1 to 10^-20^) and (B) it was statistically significant for e-values in the range from 1 to 10^-10^. Odd ratios are shown on the x-axis (log-scale). Solid dots mark contingency tables where we detected statistically significant bias by Fisher’s exact test (*p* < 0.05). Error bars represent 95% confidence interval. More details of this bias for e-value = 10^-3^ are shown in the Figure 5C.

**Supplementary Figure S5.**
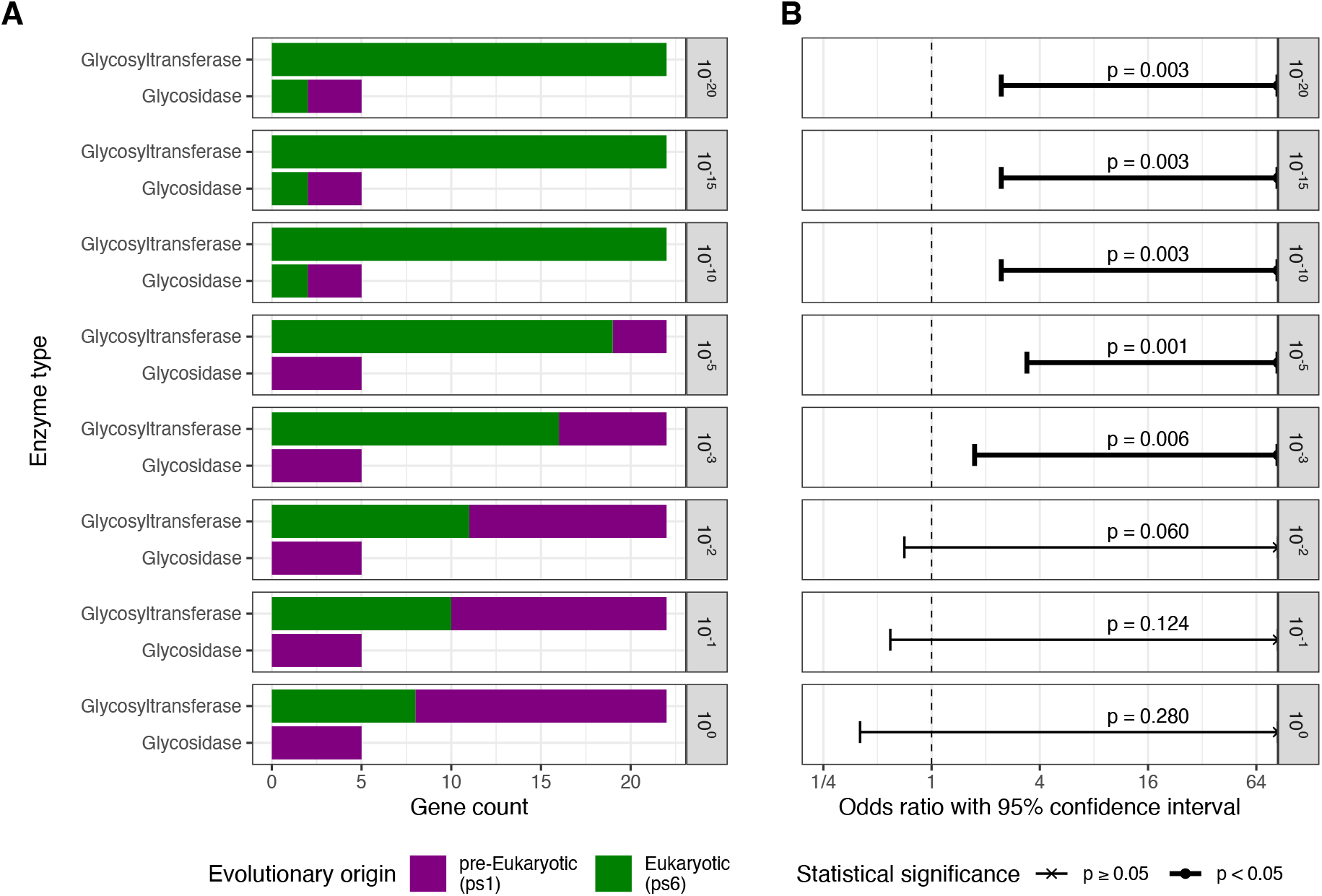
Testing the robustness of biases in the contingency table by sliding e-value analysis (Figure 6B). We detected statistically significant bias in contingency tables that compare evolutionary origin and enzyme type of N-glycosylation genes in the Golgi apparatus. (A) Glycosidases tend to be evolutionary older (Cellular organisms, ps1) compared to glycosyltransferases located in Golgi which tend to be evolutionary younger (Eukaryota, ps6). This bias was present in the full range of tested e-values (from 1 to 10^-20^), and it was statistically significant in the range between 10^-3^ and 10^-20^. Odd ratios are shown on the x-axis (log-scale). Solid dots mark contingency tables where we detected statistically significant bias by Fisher’s exact test (*p* < 0.05). Error bars represent 95% confidence interval. More details of this bias for e-value = 10^-3^ are shown in the Figure 6B.

**Supplementary Figure S6.**
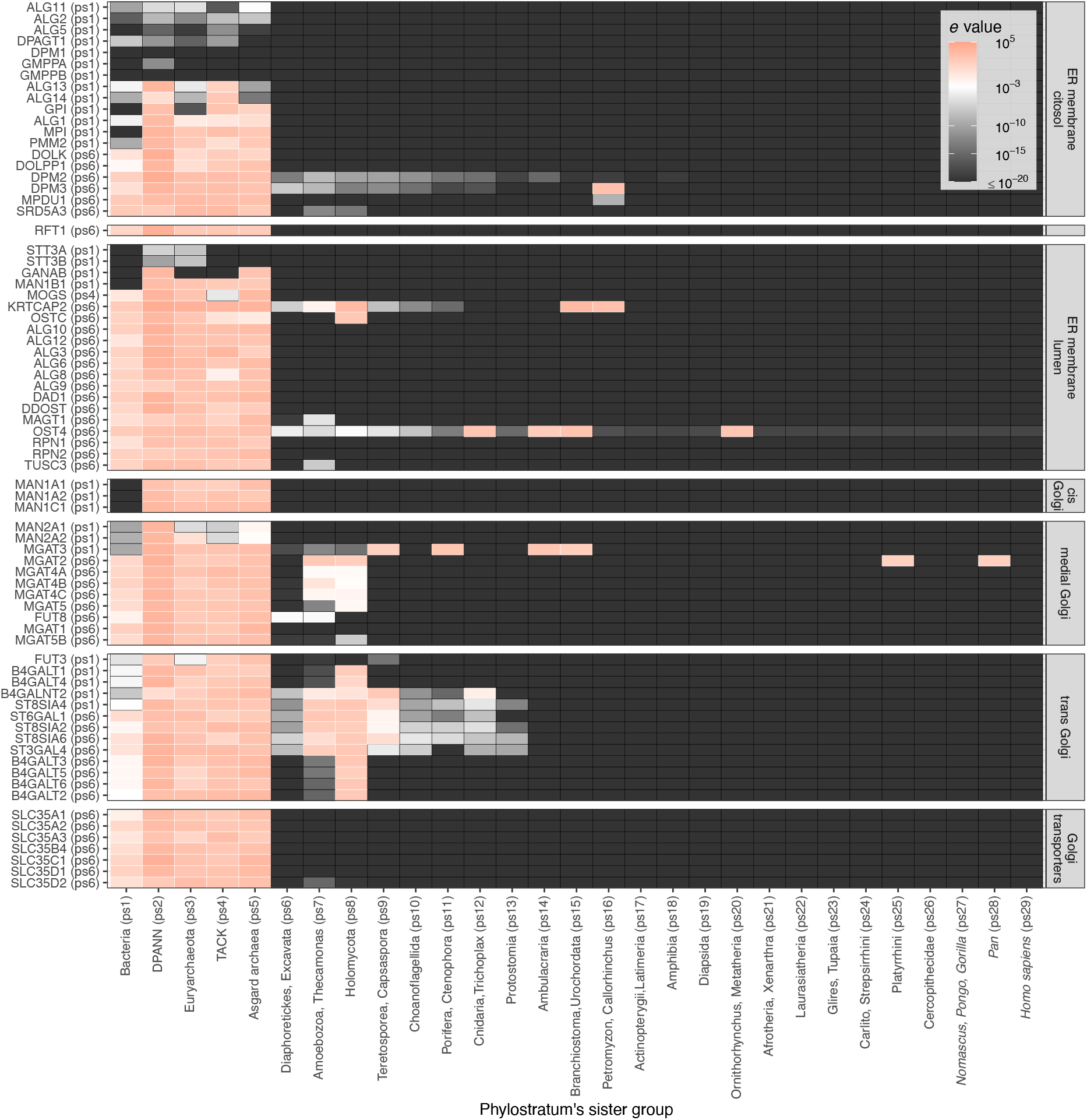
The heatmap of the best matches (minimal e-values) per phylostratum for N-glycosylation genes shown in Figure 5 and Figure 6. The matches with e-value = 10^-3^ are represented by white color, smaller e-values (better hits) by shades of gray and black rectangles, while higher e-values (weaker hits) are represented by shades of red. Gene names and corresponding phylostrata are on the y-axis. Phylostrata numbers and the names of the side branches are shown on the x-axis.

## Notes

### Competing Interest Statement

The authors have declared no competing interest.

